# Multivariate pattern analysis reveals location specific aftereffects of 10Hz motor cortex transcranial alternating current stimulation

**DOI:** 10.1101/2021.09.30.462549

**Authors:** Elinor Tzvi, Jalal Alizadeh, Christine Schubert, Joseph Classen

**Author notes:** <u>Corresponding author</u> Dr. Elinor Tzvi, Dept. of Neurology, Leipzig University, Liebigstraße 20, 04103 Leipzig, Germany.

## Abstract

**Background:** Transcranial alternating current stimulation (tACS) may induce frequency-specific aftereffects on brain oscillations in the stimulated location, which could serve as evidence for region-specific neuroplasticity. Aftereffects of tACS on the motor system remain unknown.

**Objective:** To find evidence for aftereffects in short EEG segments following tACS to two critical nodes of the motor network, namely, left motor cortex (lMC) and right cerebellum (rCB). We hypothesized that aftereffects of lMC will be stronger in and around lMC compared to both active stimulation of rCB, as well as inactive (“sham”) control conditions.

**Methods:** To this end, we employed multivariate pattern analysis (MVPA), and trained a classifier to distinguish between EEG signals following each of the three stimulation protocols. This method accounts for the multitude facets of the EEG signal and thus is more sensitive to subtle modulation of the EEG signal.

**Results:** EEG signals in both theta (θ, 4-8Hz) and alpha (α, 8-13Hz) were better classified to lMC-tACS compared to rCB-tACS/sham, in and around lMC-tACS stimulation locations (electrodes FC3 and CP3). This effect was associated with a decrease in power following tACS. Source reconstruction revealed significant differences in premotor cortex but not in primary motor cortex as the computational model suggested. Correlation between classification accuracies in θ and α in lMC-tACS was stronger compared to rCB-tACS/sham, suggesting cross-frequency effects of tACS. Nonetheless, θ/α phase-coupling did not differ between stimulation protocols.

**Conclusions:** Successful classification of EEG signals to left motor cortex using MVPA revealed focal tACS aftereffects on the motor cortex, indicative of region-specific neuroplasticity.

## Introduction

Transcranial alternating current stimulation (tACS) has been previously shown to modulate endogenous oscillations in a frequency-specific manner and a range of behaviors [1]. TACS affects endogenous oscillations “online”, during the stimulation, and “offline”, referring to effects that persist or arise after the stimulation has been terminated. It has been hypothesized that “online” effects occur through entrainment, i.e., alignment of brain oscillations to the periodic external signal by tACS. In contrast, neuroplasticity is the dominant theory regarding “offline” effects occurring at the target location, possibly through spike-timing dependent plasticity (STDP).

The “offline” aftereffect has been explored experimentally by applying tACS at 10Hz or the individual α frequency (iAF) over parieto-occipito areas during rest [2,3], as well as during task performance [4–7]. These studies found posterior α power increase in post-compared to pre-stimulation blocks, during either rest or task performance. However, over the sensorimotor cortex, an opposite effect was demonstrated: iAF-tACS applied over centro-parietal areas during rest led to a decrease in µ power, i.e., 8-12Hz oscillatory power over the motor cortex. Thus, it seems that α-tACS induces topographically heterogeneous aftereffects within the motor network that depend on the stimulation site. However, it remains unclear whether tACS may induce specific effects on the motor network when specific nodes of the network are targeted.

Here, we aimed to investigate aftereffects on the motor network induced by tACS of left motor cortex (lMC) by re-analyzing a previously published data-set [8]. Results were compared to an active (tACS to right cerebellum) and an inactive (“sham” stimulation) control conditions. To this end, we employed multivariate pattern analysis (MVPA), a method that extends beyond standard univariate techniques by exploiting the interactions between multiple features of the EEG signal, such as spectral profiles across multiple channels or sources, using machine-learning algorithms [9]. MVPA has an advantage over classic parameter estimation in that it allows data-driven analyses with few a-priori hypotheses regarding spatial or temporal patterns. Here, we tested whether a classifier is able to discriminate oscillatory activity recorded after tACS has terminated, at single-trial level, between specific tACS protocols. Thus, it provides a means to statistically assess differences in oscillatory patterns between stimulation protocols.

We hypothesized that tACS-aftereffects are: (1) most unequivocally detected at or near the location of the stimulation, (2) specific to the stimulation frequency, (3) evident on single-trial data, and (4) during a visuomotor task as well as rest.

## 1. Methods

### 1.1. Participants and Experimental design

Details on participant inclusion and experimental design can be found in [8]. In short, 25 young (mean age: 24.8 years) healthy participants received 10Hz tACS to either left motor cortex (lMC) or right cerebellum (rCB) as well as sham stimulation in alternate to either location (Fig. 1), in separate experimental sessions at least one week apart. One participant was excluded due to a technical error in the data acquisition resulting in a total sample of 24 participants. During stimulation, participants learned a motor sequence (see details in supp. materials, section 1). Before and after tACS, 64-channel EEG was collected in two conditions: resting-state with eyes-open (RSEO) or eyes-closed (RSEC), each with a duration of 200s, or during a simple stimulus-response matching task (see detailed description in supp. materials, section 1).

**Figure 1.**
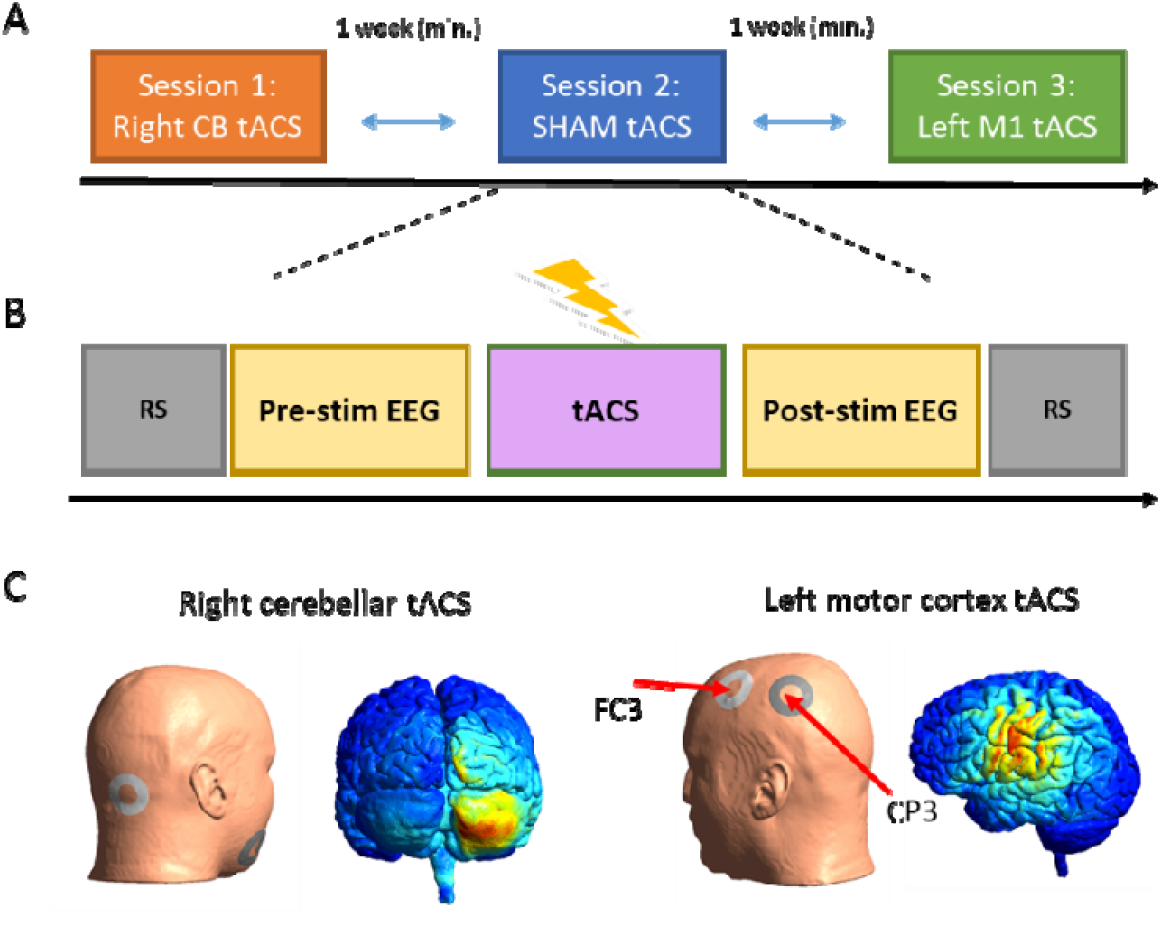
Experimental design. **A** Experimental sessions were kept at least a week apart and were counter-balanced between subjects. **B** Each session was divided into a pre-stimulation and post-stimulation segments, each with a resting-state recordings (both eyes-closed and eyes-open, marked in grey) as well as in a stimulus-response task (marked in yellow). **C** computational modelling of electric field distribution in left MC and right cerebellum during tACS.

### 1.2. Transcranial alternating current stimulation (tACS) protocol

Transcranial alternating current stimulation (tACS) was applied (DC-Stimulator PLUS, NeuroConn, Ilmenau, Germany) via two ring-shaped conductive rubber electrodes with an outer diameter of 48mm, and an inner diameter of 24mm (area: 15cm^2^) and an intensity of 1mA at 10Hz (peak-to-peak-amplitude; sinusoidal waveform; 0.07 mA/cm^2^ current density) for a total duration of 20min. For rCB-tACS, one electrode was placed on the right mandibula and the other 1cm below and 3cm right to the inion. For lMC-tACS, one ring-shaped electrode was placed around electrode FC3 and one around CP3 rendering the current flow as precisely as possible to C3 (Fig. 1C) as shown by computational simulations (see supp. materials, section 2). For sham stimulation, the current was ramped up for 30s, then stayed at 1mA for 10s and ramped down for another 30s, in order to effectively blind the participants to the experiment protocol. Awareness to the stimulation protocol was evaluated using a questionnaire (see supp. materials, section 3, for details).

### 1.3. EEG recordings and pre-processing

EEG was recorded using Ag/AgCl electrodes embedded in a 64-channel cap and connected to an eegoTM amplifier (ANT Neuro b.v., Hengelo, the Netherlands) with a sampling rate of 512Hz and 24bit resolution. A low-pass filter was applied at 0.26*sampling rate (f_c ≈ 133Hz). Eye movements were recorded with an electrooculogram below the left eye. EEG was recorded against an online reference electrode in location CPz. Pre-processing and all subsequent analyses were performed using in-house MATLAB® (The MathWorks, Natick, MA, USA) scripts and the EEGLAB toolbox [10]. Signals were band-pass filtered (F_*cutoff*_ = 1 - 49Hz) to remove slow drifts and power line noise and re-referenced offline to the average of the signal from left and right mastoids. The signal from electrode CPz was re-calculated. Next the signals were segmented into 3s epochs for the task-based data and 4s (non-overlapping) epochs for the RSEO and RSEC data. Using ICA, 3-4 components related to eye blink artifacts were identified and removed. Additional artifacts were removed using a simple threshold (−70µV, +70µV) on the filtered data and finally, the signals were re-referenced to a common-average reference.

### 1.4. EEG spectral power and multivariate pattern analysis

We used the Fieldtrip integrated MVPA-light toolbox [11] to perform multivariate pattern analysis (MVPA), using regularized multi-class Linear Discriminant Analysis classifier. In Fig. 2, we provide an overview of the analysis pipeline for both scalp and source-based signals. First, we computed the power spectrum of EEG signals in a 400ms time window using a Morlet wavelet. Signals were filtered to obtain oscillatory power at 1–49Hz using wavelets of 7 cycle length. Frequency resolution was set to 1Hz and time resolution to 10ms. Then we calculated the spectral power difference before (PRE) and (POST) after tACS. Each of the data segments (PRE, POST) contained ∼80 epochs for task-based data and ∼170 epochs for RSEO and RSEC data. To calculate POST-PRE power difference (ΔPOST-PRE), we randomly shuffled the order of the epochs in PRE and in POST. ΔPOST-PRE power for each shuffled trial pairs, each time-point and each electrode, was normally distributed and used as an input for the classifier. Classification was performed at each frequency component in 4-49Hz range separately. We specified the time domain as the feature dimension for the classifier. This means that the classifier searched for the optimal time segments within the specified time window (see above) for the classification process. The performance of the classifier was then evaluated for each electrode.

**Figure 2.**
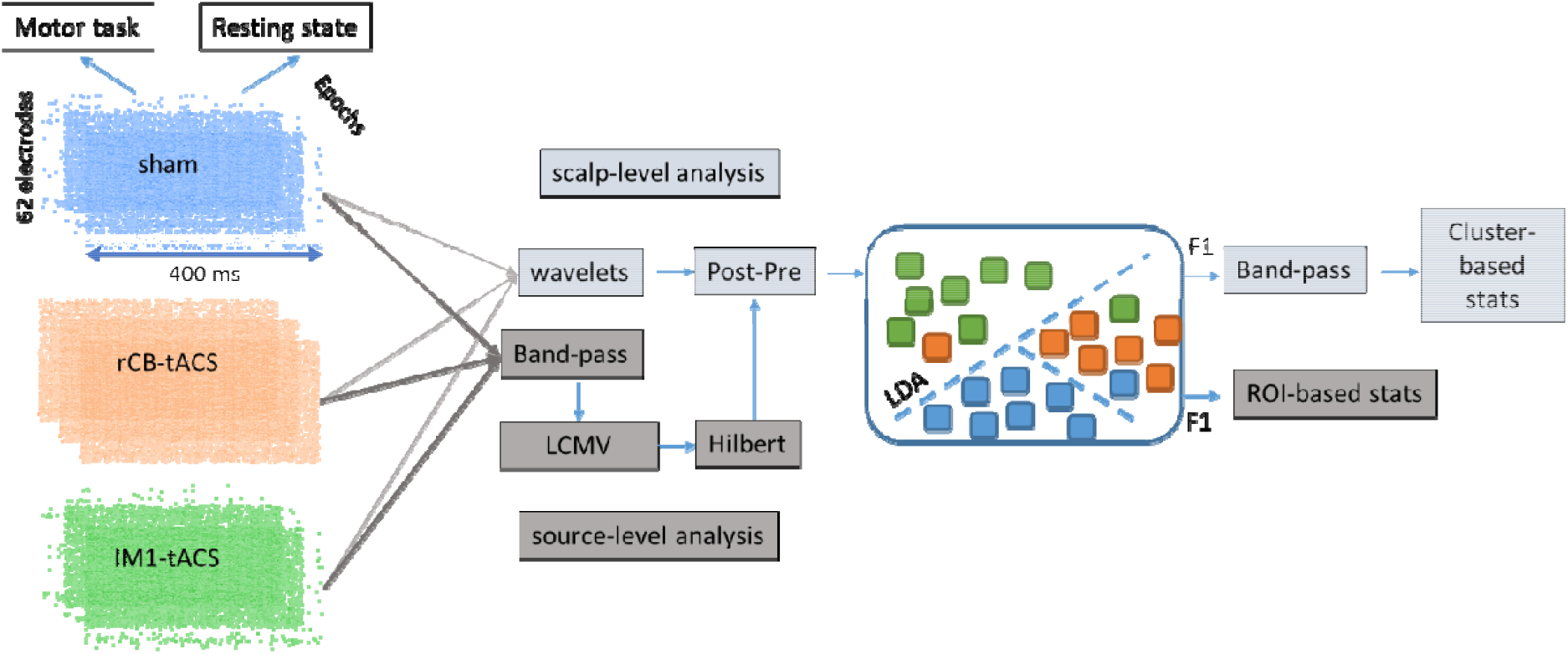
Analysis pipeline. Short EEG epochs in each stimulation protocol were analyzed on both scalp level (using electrodes, light grey) and source-level (using reconstructed voxels, dark grey). Post-Pre power differences were then analyzed using a multi-class LDA which produced a classification accuracy (F1-score) for each stimulation protocol. Differences in accuracies were analyzed on scalp-level using cluster-based statistics, and on source-level using a ROI-based approach.

For classification, data were first projected into 2-dimensional discriminative subspace. Then, an optimal linear transformation of the data was applied to maximize class separability. To counteract over-fitting, a regularization parameter λ was estimated. The procedure entailed: training, testing and cross-validation. During training, the parameters of the model were optimized to discriminate between the three classes (lMC-tACS, rCB-tACS and sham). During testing, the model performance was evaluated on an independent subset of the data which was not used for training. The training and testing phases were repeated on different subsets of the data using 10-fold cross-validation, in which the data was split into 10 different parts and in each iteration of the cross-validation phase, one of the 10 parts was used for testing and the other 9 parts for training. The performance of the classifier was evaluated using the F1-score, which is the harmonic average of precision and recall.

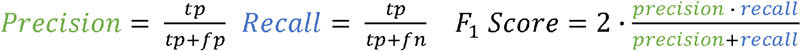

Where tp is the number of true positives, fp - the number of false positives, and fn - the number of false negatives. Importantly, for multiclass classification, F1-score serves as the most optimal score for evaluating classification accuracy. Finally, the F1-score was averaged across theta (θ, 4–8Hz), alpha (α, 9-13Hz), beta (β, 14-30Hz) and gamma (γ, 31-49Hz).

### 1.5. Source reconstruction and multivariate pattern analysis

To reconstruct the EEG signals in source space we used the linear constrained minimum variance (LCMV) beamforming approach. Details on the reconstruction procedure are found in the supp. Materials (section 4). Following the pre-processing steps described above, signals were band-pass filtered for α and θ, in accordance with the scalp-level results below. Next, the Hilbert transform was used to obtain the spectral power. The MVPA procedure was then performed identically to the electrode-space analysis above, only that instead of electrodes, the performance of the classifier was evaluated for each grid point. In order to prevent spurious classification results due to spatial leakage [12], we used a parcellation procedure based on an automated Talairach atlas [13] as well as the AAL (automated anatomical atlas) [14] to create regions of interest (ROIs). F1-scores were averaged across grid points belonging to each ROI. Based on the scalp-level results, we specified five ROIs on the left hemisphere: M1 defined as Brodmann area (BA) 4, somatosensory cortex (S1), premotor cortex (PMC, BA6), superior parietal lobule (SPL) and inferior parietal lobule (IPL) (Fig. 6A).

### 1.6. Statistical analyses of classification accuracies

Subject-specific classification results were analyzed at the group level. To test for F1-score differences across classes (lMC-tACS, rCB-tACS and sham) in the electrode-space signals, we used a non-parametric cluster-based Monte Carlo permutation testing with 1000 randomizations as implemented in the Fieldtrip toolbox. This analysis resulted in a F-value for the comparison across all classes in each electrode and each frequency band. For the source-space data, we tested for F1-score differences across classes using the non-parametric Kruskal-Wallis test in each of the five ROIs (M1, S1, PMC, SPL and IPL). P-level threshold was FDR-corrected across the five ROIs. All post-hoc tests of significant effects were performed using the non-parametric Wilcoxon signed-rank test.

## 2. Results

### 2.1. Better classification of α and θ power to lMC tACS is specific to stimulation electrodes FC3 and CP3

#### Task-based analysis

F1-score of each class was subjected to a 1-way rmANOVA and cluster-based permutation analysis of each electrode in each frequency band. We compared lMC-tACS aftereffects to both rCB-tACS and sham. We hypothesized that successful classification of spectral α power to lMC-tACS would be significantly larger in electrodes at and adjacent to the stimulation electrodes FC3 and CP3, when compared to both rCB tACS and sham. Indeed, for α, we found a left central cluster (p = 0.047) with maximal effects in CP1 (F_2,23_ = 6.5) but also CP3 (F_2,23_ = 5.7) and FC3 (F_2,23_ = 5.1) (see Fig. 3A). For θ, a similar cluster was observed (cluster p = 0.016) with maximal effects at FC3 (F_2,23_ = 8.9) and CP3 (F_2,23_ = 7.4) (Fig. 3B). No significant clusters were observed for β or γ. In Fig. 3C, we plot the mean F1-score for each frequency component in each of the electrodes of the cluster identified above.

**Figure 3.**
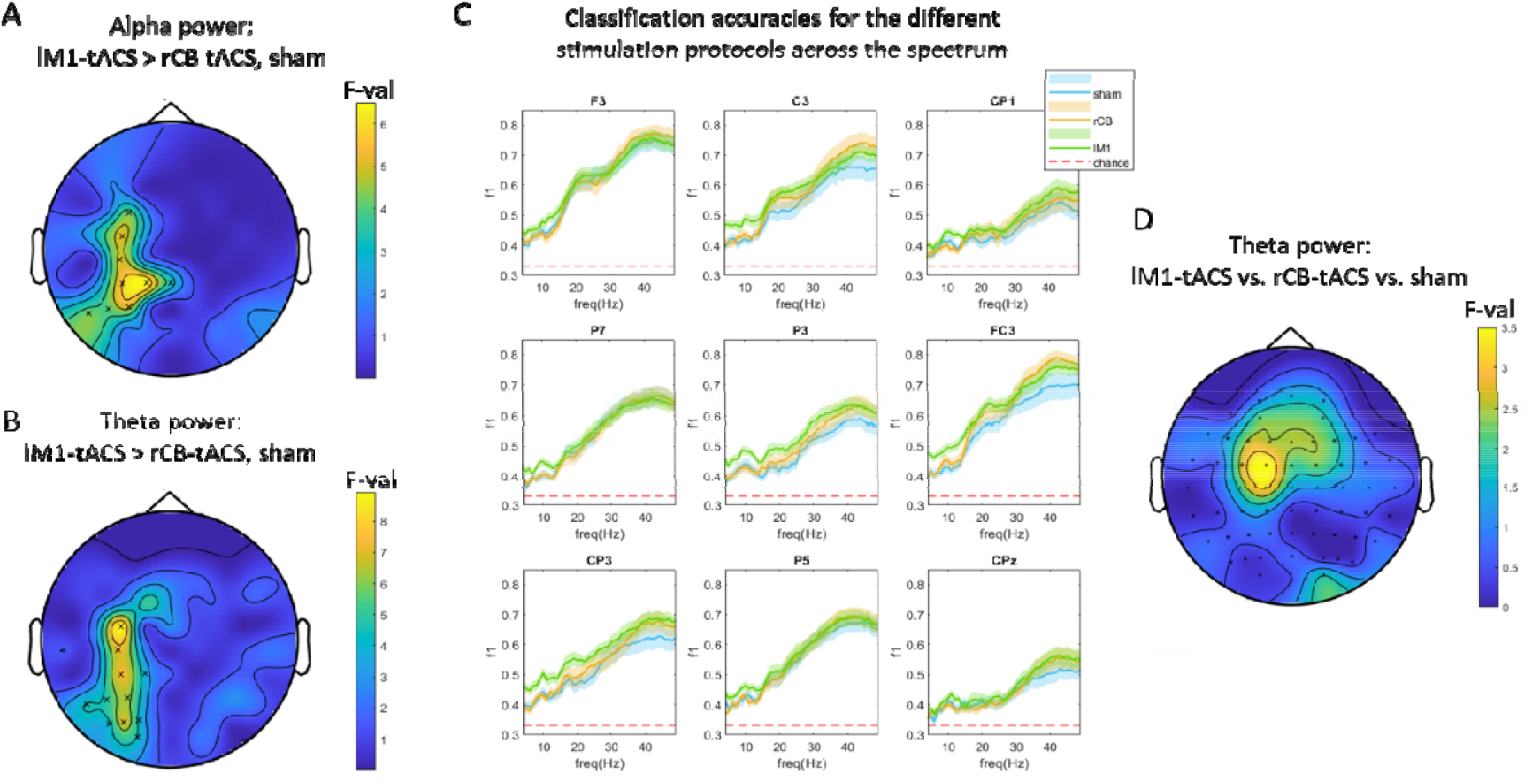
Task classification results. **A-B** topographic plots of F-values showing differences between classification accuracies of POST-PRE, single-trial, alpha (A) and theta (B) power across stimulation protocols. **C** Spectral representation of classification accuracies for each stimulation protocol in electrodes of the cluster shown in A. Shaded areas represents the standard error of the mean across subjects. **D** Topographic plot for F-values showing theta POST-PRE power differences (uncorrecred).

Post-hoc signed rank tests on classification effects revealed that the significant difference between the three classes in α stems from larger classification accuracy for lMC-tACS compared to rCB-tACS in most electrodes of this cluster (Fig. 3A, C, Z > 2.1, p < 0.05, FDR corrected). Classification accuracy comparing lMC-tACS to rCB-tACS in θ showed significant differences in electrodes C3, CP1, P7, P3, O1, FC3, CP3, P5, PO5, PO3 and PO7 (all Z > 2.1, p < 0.05, FDR corrected).

Comparing classification accuracy in lMC-tACS to sham, significant difference in θ were found for FC3 only (Z = 2.9, p = 0.003, FDR corrected). A similar difference in CP3 was found on trend (Z = 2.4, p = 0.016, no correction). For α, a similar trend (p < 0.05, no correction) was found in electrodes FC3 and CP3 (Z = 2.7, p = 0.008; Z = 2.3, p = 0.02).

*In sum, these results demonstrate a location specific aftereffect of 10Hz tACS to the lMC on single-trial* α *and* θ *power, during performance of a motor task*.

#### Resting-state analysis

We performed a similar analysis in resting-state eyes-open (RSEO) as well as resting-state eyes-closed (RSEC) of ΔPOST-PRE power. In accordance with the task-based signals, a significant left-central cluster was evident in α (p = 0.016) for RSEO. The cluster for RSEO had maximal effects in electrode P3 (F_2,23_ = 12.2, see topoplot in Fig. 4A), adjacent to the stimulation electrode CP3. A strong effect was found also in FC3 (F_2,23_ = 6.5). For θ, a similar cluster was also found (Fig. 4B, cluster significance: p = 0.04) with a maximal effect in FC3 (F_2,23_ = 12.9). Fig. 4C shows the mean F1-score for each frequency component. No clusters were evident for RSEC.

**Figure 4.**
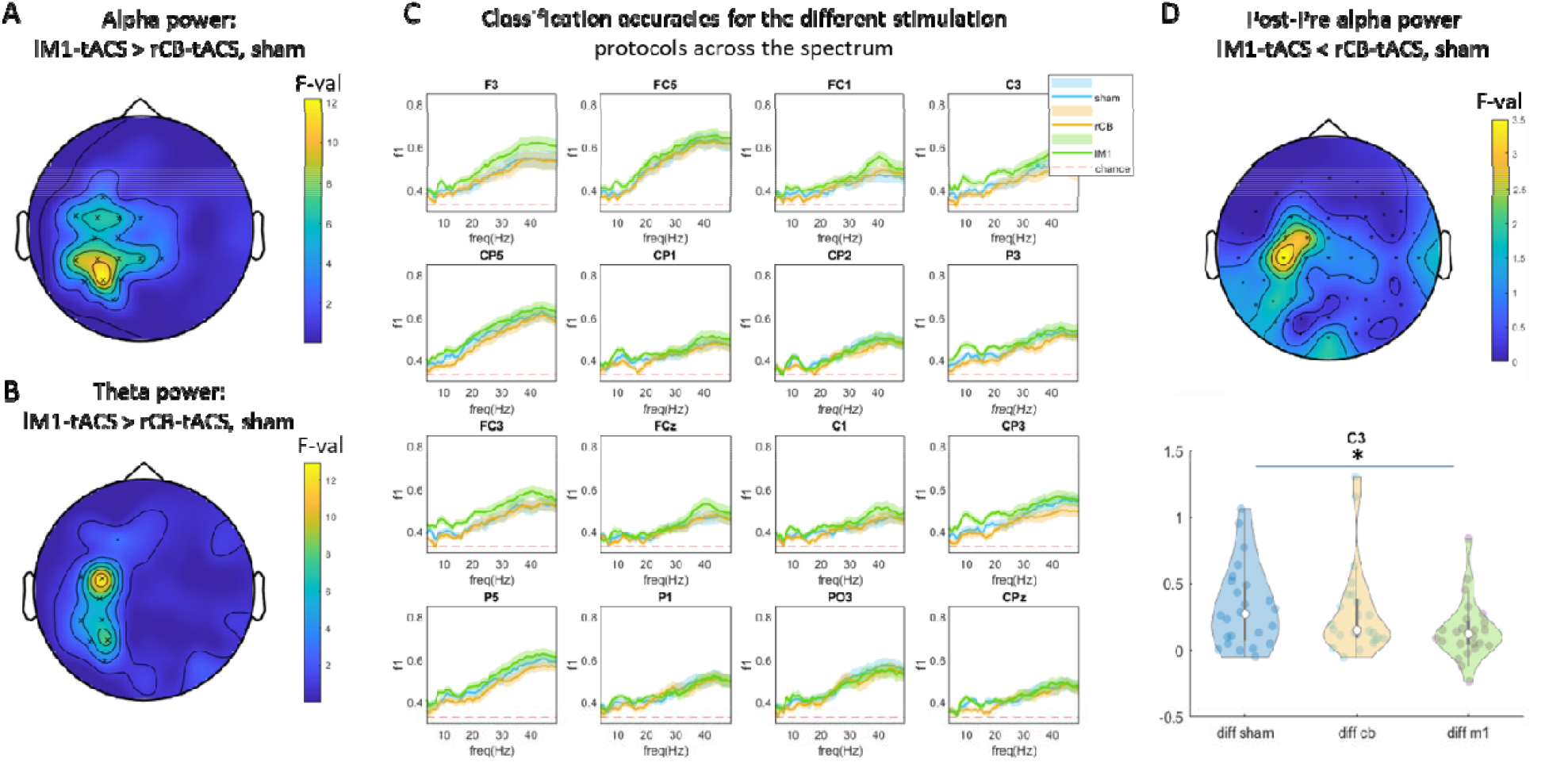
Resting-state classification results. **A-B** topographic plots of F-values showing differences between classification accuracies of POST-PRE, single-trial, alpha (A) and beta (B) power across stimulation protocols. C Spectral representation of classification accuracy for each stimulation protocol in electrodes of the cluster shown in A. Shaded areas represents the standard error of the mean across subjects. D Topographic plot for alpha POST-PRE power differences. In electrode C3 POST-PRE power was lowest for lM1-tACS compared to both rCB-tACS and sham (violin plot).

Post-hoc signed rank tests on classification effects in α revealed that the significant difference between the three classes stemmed from larger classification accuracy for lMC-tACS compared to rCB-tACS in the entire cluster (Fig. 4A, 4C, all Z > 2.2, p < 0.05, FDR corrected). For θ, classification accuracy was larger for lMC-tACS compared to rCB-tACS as well in the entire cluster (Fig. 4B, all Z > 2.2, p < 0.05, FDR corrected). Compared to sham, accuracy for lMC-tACS in α was significantly larger in electrodes CP3 (Z = 3.1, p = 0.002) and P3 (Z = 3.4, p < 0.001). For θ, accuracy for lMC-tACS tended to be larger compared to sham in electrodes P3 (Z = 2.2, p = 0.028) and FC3 (Z = 2.4, p = 0.018).

*These results further corroborate a location specific aftereffect of 10Hz tACS to lMC on single-trial* α *and* θ *power of eyes-open resting-state signals*.

### The effect of tACS on θ/α power differences

Next, we averaged oscillatory power across all trials during task performance in PRE and POST tACS blocks, in order to determine the direction of power changes following tACS. Then, we assessed ΔPOST-PRE power between the stimulation protocols using whole-brain Monte-Carlo permutation analysis. There were no significant differences between the stimulation protocols on a corrected level (cluster based, p < 0.05). However, electrodes FC1 (F_2,23_ = 3.7, p = 0.02) and C1 (F_2,23_ = 3.2, p = 0.037) showed a tendency for a significant difference across stimulation protocols in θ (Fig. 3D). Post-hoc Wilcoxon signed rank tests showed no significant differences. There were no differences evident for α.

A similar analysis of RSEO signals revealed no significant differences between stimulation protocols on a corrected level (cluster-based p < 0.05), but electrode C3 showed evidence for differences in α power (F_2,23_ = 3.7, p = 0.02, Fig. 4D). Indeed, post-hoc Wilcoxon signed rank tests showed that ΔPOST-PRE power under lMC-tACS were smaller compared to both rCB-tACS (Z = 2.3, p = 0.02) and sham (Z = 2.0, p = 0.045). No such effects were observed for θ. There was no correlation between ΔPOST-PRE power and F1-scores in any of the classes.

*These results show that (1) traditional analysis of power effects averaged across trials are far less sensitive compared to MVPA. (2) classification differences between lMC-tACS and rCB-tACS/sham result from power decrease post-stimulation*.

### Association between aftereffects of tACS on θ and α frequency bands

We further explored whether classification accuracy in electrodes FC3, C3 and CP3 in α was associated with classification accuracy in θ, which would raise the possibility that aftereffects in θ were driven by the narrow band effects around the stimulation frequency. Indeed, in the task-based signals, F1-scores at θ were correlated with F1-scores at α for lMC-tACS in FC3, C3 and CP3 (r = 0.85, 0.79, 0.65, all p < 0.001, Fig. 5A). Using Fisher Z-transformation, we found that correlation between F1-scores in θ/α in electrode FC3 was significantly stronger in lMC-tACS compared to rCB-tACS (p = 0.03) and sham (p = 0.01). Similarly, a correlation analysis between classification accuracies of α and θ for RSEO signals showed a strong association in electrodes FC3, C3 and CP3 for lMC-tACS (r = 0.66, 0.66, 0.68, p < 0.001, see Fig. 5A). Here however, the correlation coefficients did not differ between lMC-tACS and rCB-tACS/sham in any of the electrodes (p > 0.06, uncorrected).

**Figure 5.**
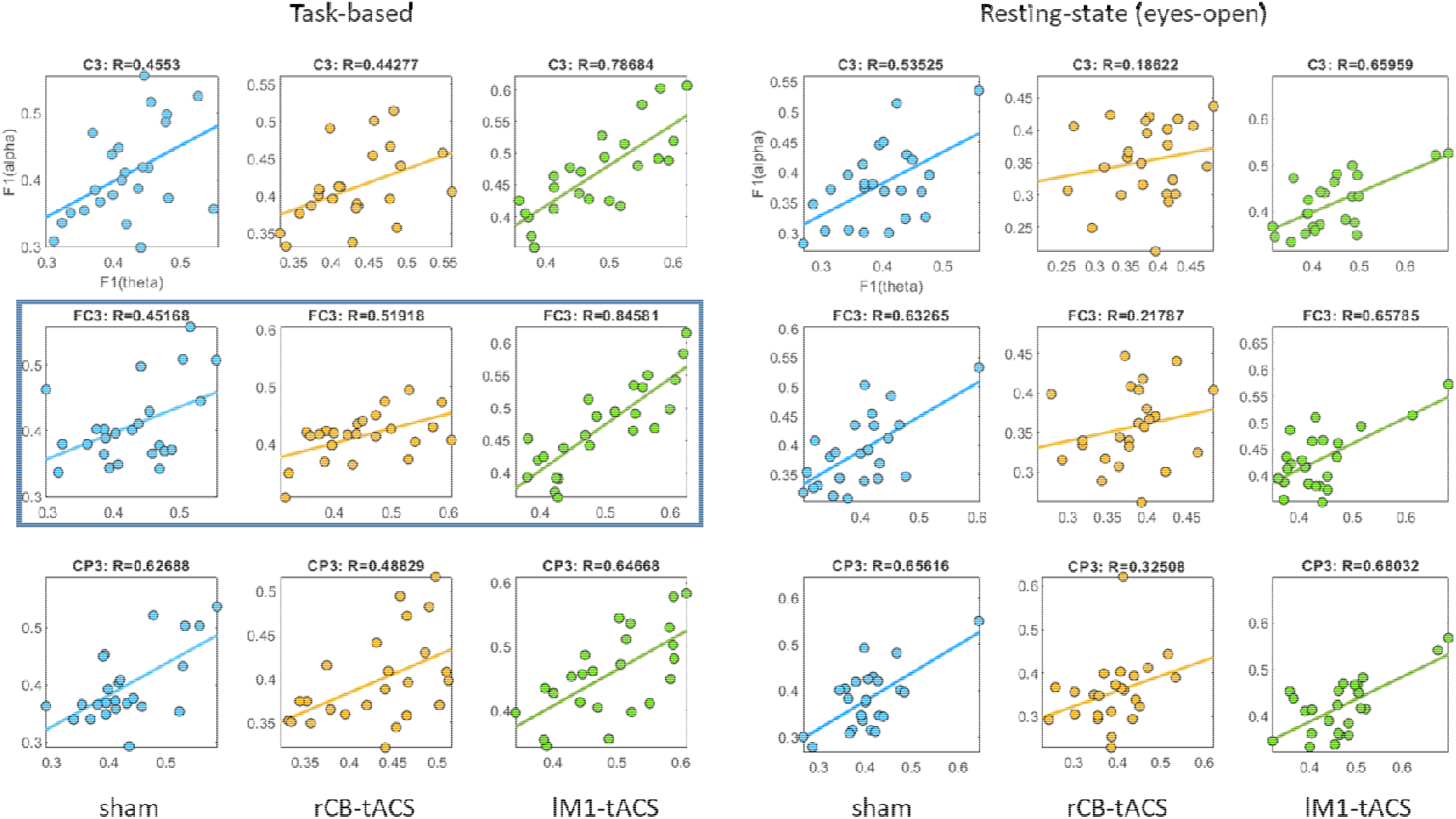
Correlation between theta and alpha classification results. F1-score for theta and alpha frequency band was strongly correlated for lM1-tACS in both task-based (A) and resting-state signals (B). In electrode FC3 and task-based signals (marked with a frame) this correlation in was larger compared to both rCB tACS and sham

To tap into possible mechanisms of these associations, we further explored θ/α phase-coupling in electrodes FC3, C3, and CP3, averaged across all trials in PRE and POST tACS blocks (see supp. Materials, section 5, for a description of this analysis). We found no differences in phase-coupling between stimulation protocols for task-based signals (all p > 0.4). For RSEO signals, significant differences between the stimulation protocols were found in FC3 (p = 0.01, data not shown) but post-hoc Wilcoxon signed-rank tests showed that this effect stemmed from increased θ/α phase-coupling in rCB-tACS compared to sham (Z = 2.7, p = 0.006) and a tendency for larger phase-coupling in lMC-tACS compared to sham (Z = 1.7, p = 0.08). These results suggest that the stronger associations between classification accuracies in θ and α under lMC-tACS are likely not driven by phase-coupling mechanisms.

*Together, these results suggest that localized aftereffects of 10Hz lMC-tACS were broad band and not frequency specific*.

### Localization of classification effects in source-space

To further study the spatial topography of better classification of ΔPOST-PRE in α and θ power to lMC-tACS compared to rCB-tACS and sham, we used LCMV beamforming to reconstruct classification accuracies in source-space. We focused our analysis on the following left hemispheric ROIs: M1, S1, PMC, SPL and IPL, based on the scalp-level results (Fig. 6A). First, as classification accuracies at source-level were lower compared to scalp-level, we evaluated in each ROI whether classification was better compared to chance level (approximated at 34% across subjects and classes, FDR corrected for multiple comparisons across all ROIs).

**Figure 6.**
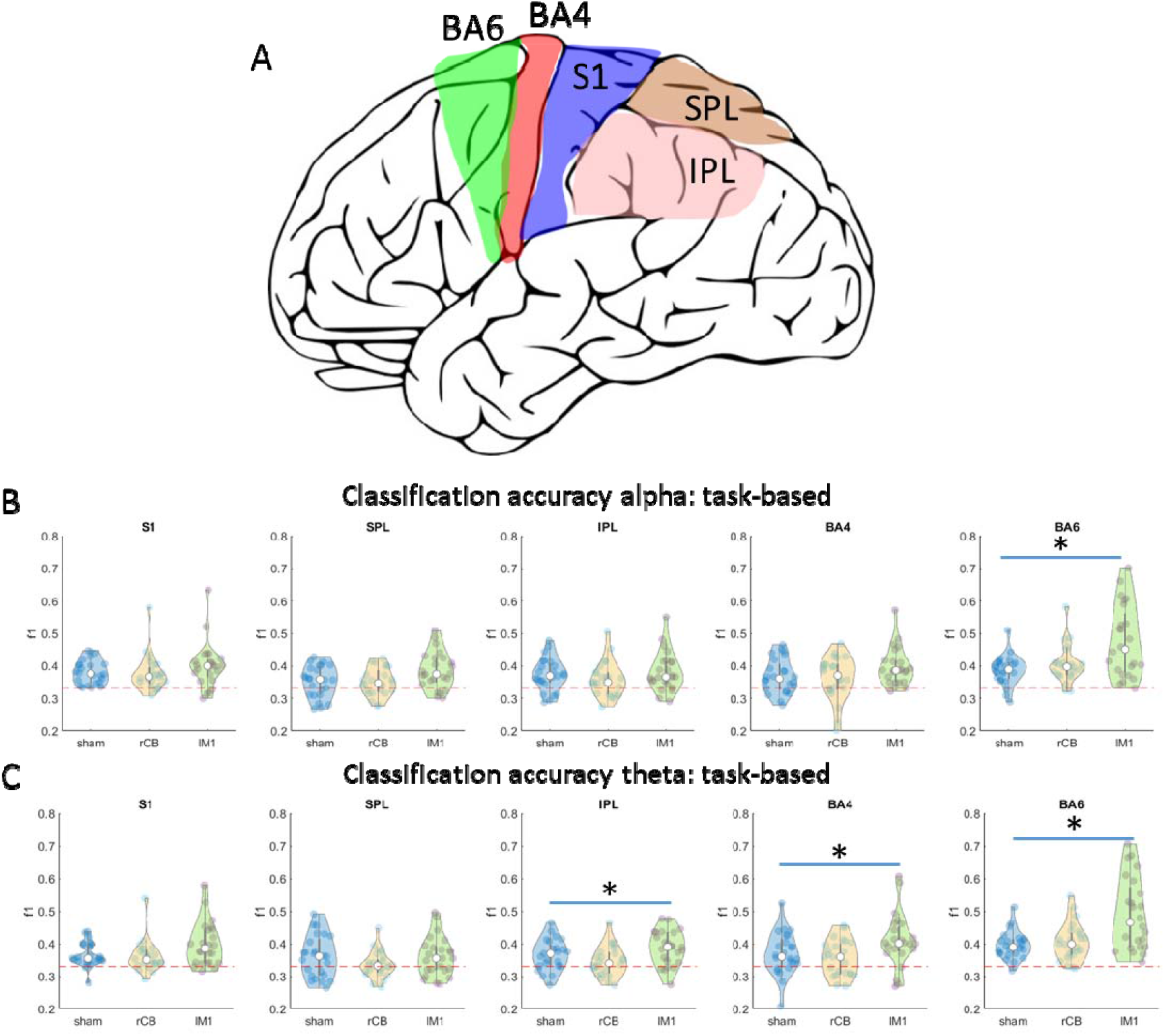
Classification of task-based signals in source space. **A** illustration of regions-of-interest included in the source-space analysis. **B-C** classification accuracies averaged across all voxels in somatosensory cortex (S1), primary motor cortex (BA4), premotor cortex (BA6), superior parietal lobule (SPL) and inferior parietal lobule (IPL) in alpha (**B**) and theta (**C**).

**Figure 7.**
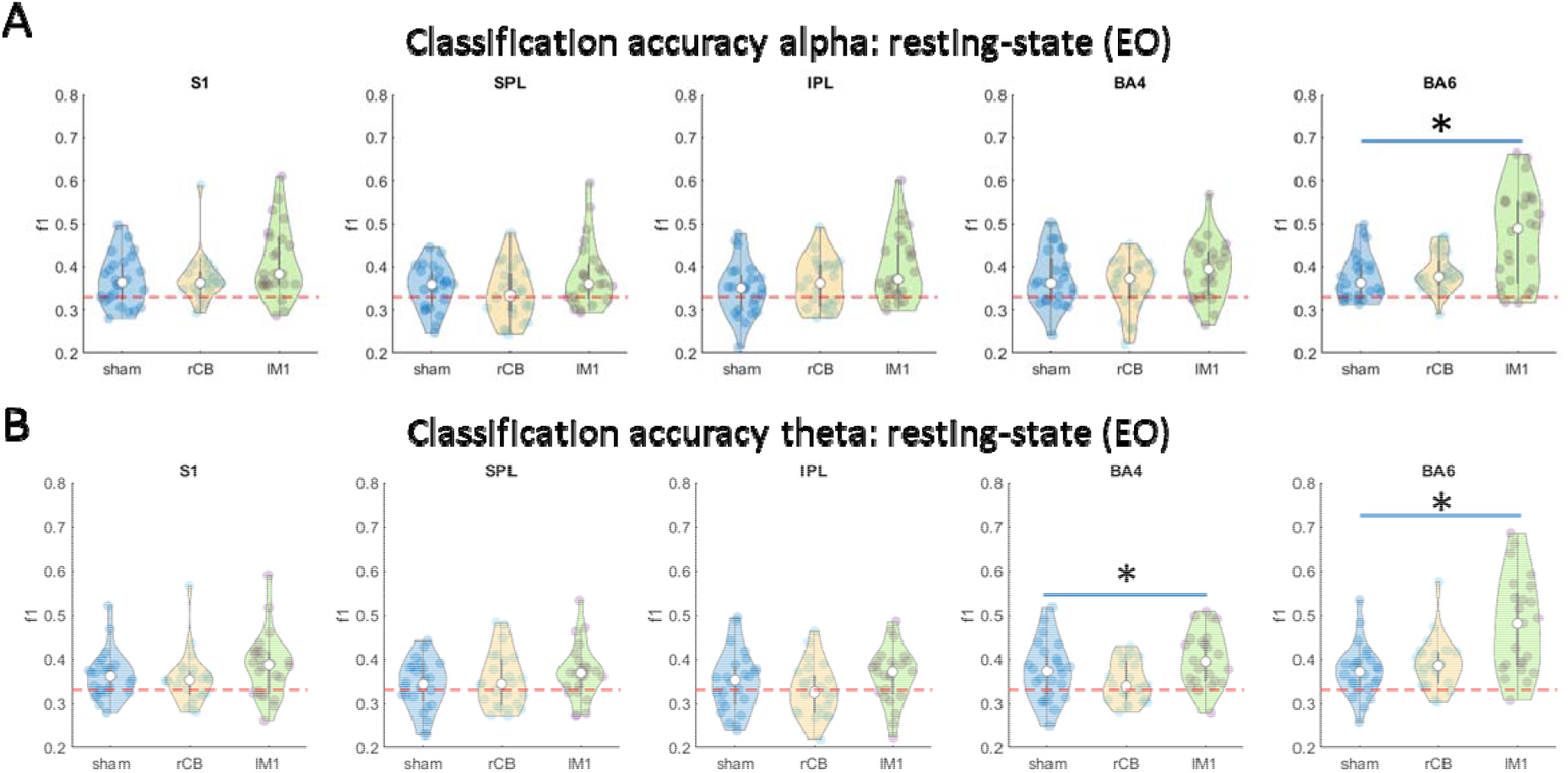
Classification resting-state signals in source space for alpha (A) and theta (B).

#### Task-based signals

Across all classes, classification accuracies of reconstructed α power in ROIs: M1, S1, PMC and SPL were significantly better than chance level. For reconstructed θ power, classification accuracies in S1 and PMC were better than chance level across all classes.

We then compared classification accuracies of α and θ power across the three classes in each ROI. For α, we found a significant effect in PMC (χ^2^ = 10.3, p = 0.006, Fig. 6B). Post-hoc tests revealed that this effect in PMC stemmed from a larger classification accuracy in lMC tACS compared to sham (Z = 3.0, p = 0.003) as well as compared to rCB tACS (Z = 2.3, p = 0.02). For θ (Fig. 6C), a significant effect was found in IPL (χ^2^ = 8.3, p = 0.016), M1 (χ^2^ = 8.5, p = 0.014) and PMC (χ^2^ = 11.5, p = 0.003). Post-hoc tests revealed that these effects stemmed from significant difference in lMC-tACS compared to sham in PMC (Z = 3.1, p = 0.002). In M1, the effects stemmed from significant difference in lMC-tACS compared to rCB-tACS (Z = 2.9, p = 0.004).

#### Resting state

Classification accuracy of reconstructed α power in RSEO was better than chance level across all classes in ROIs: S1, M1 and PMC. Significant difference in classification accuracies across classes was observed only in PMC (χ^2^ = 10.7, p = 0.005, Fig. 6D). Post-hoc tests revealed that these effects stemmed from larger classification accuracy in lMC-tACS compared to sham (Z = 2.8, p = 0.005) and rCB-tACS (Z = 2.3, p = 0.02). Classification accuracy of reconstructed θ power in RSEO was better than chance level across all classes only in PMC. Significant differences in classification accuracies across classes were observed in M1 (χ^2^ = 8.3, p = 0.02) and PMC (χ^2^ = 13.3, p = 0.001) (Fig. 6E). Post-hoc tests in PMC revealed larger classification accuracy in lMC-tACS compared to sham (Z = 3.2, p = 0.002) and rCB-tACS (Z = 2.5, p = 0.01).

*These results suggest that better classification of* θ *and* α *single-trial power to lMC-tACS observed in stimulation electrodes FC3 and CP3 on the scalp-level, was based on a source residing in left PMC*.

## Discussion

In this study, we examined the aftereffects of 10Hz tACS on key nodes of the motor network using multivariate pattern analysis (MVPA). Our results show that tACS aftereffects are: (1) location specific, i.e., classification to lMC-tACS was better compared to rCB-tACS/sham in electrodes FC3 and CP3 used for lMC-tACS stimulation. (2) Found in both α (9-13Hz) and θ (4-8Hz) frequency bands. (3) Evident during motor task performance as well as rest. (4) Showing specific power decrease following lMC-tACS. And (5) source localized to the premotor cortex.

To study 10Hz tACS aftereffects, as a measure of neuroplasticity, we trained a machine-learning algorithm to classify short EEG segments following tACS to the different stimulation protocols. Next, we estimated the classification accuracy of each stimulation protocol, or in other words how well the model could predict to which stimulation protocol other EEG segments, unseen by the model, belong. We then compared classification accuracies for lMC-tACS to both active rCB-tACS as well as inactive (sham) control conditions. Our results show that oscillations in both θ and α frequency bands could be better classified to lMC-tACS, specifically in and around stimulation locations (FC3 and CP3). This broad-band effect, spanning both θ and α, agrees with previous reports indicating that oscillatory power following tACS was modulated in frequency bands beyond the actual tACS frequency. For example, 5Hz tACS over the frontal cortex during non-rapid eye movement sleep, decreased oscillatory spectral power in low (0.5-4Hz) frequencies as well in the α band at the stimulation electrodes [15]. Others found increased delta power following iAF-tACS [4]. We found that classification accuracy in θ and α were correlated in FC3 and more strongly for lMC-tACS compared to rCB-tACS/sham, suggesting that these effects were strongly associated. According to Veniero et al. [16], such associations in tACS aftereffects may be rooted in cross-frequency coupling mechanisms. We found however no evidence for ΔPOST-PRE θ/α phase-coupling difference between stimulation protocols, thus it is unlikely that phase coupling explains multi-band aftereffects of 10Hz tACS to the motor cortex.

In a computational simulation of tACS aftereffects on a neural network model, incorporating spike-timing dependent plasticity (STDP) rules, long-term potentiation (LTP) of synaptic strength was induced when stimulation was at the frequency of the intrinsic oscillation. LTP was accompanied by an increase EEG spectral power [2]. In contrast, synaptic weakening occurred when stimulation was performed at a frequency slightly faster than the spontaneous frequency [6], conceivably resulting in decreased power post-stimulation. Here, we observed α power decrease (during resting-state, averaged across trials) in electrode C3 under lMC-tACS compared to rCB-tACS/sham. Thus, intrinsic frequencies below 10Hz may have led to synaptic weakening and location-specific aftereffects at both α and θ frequencies [6]. However, as previous report [17] found decreased power using iAF-tACS, the significance of small deviations between intrinsic and external frequencies remains open.

Interestingly, while both motor task and resting-state oscillatory θ/α power had stronger classification accuracy in FC3 and CP3 for lMC-tACS compared to rCB tACS/sham, only resting-state (eyes-open) α power yielded a significant location-specific decrease when data were averaged across trials. This dissociation testifies for the sensitivity of the multivariate approach in detecting subtle changes that are perhaps overlooked by traditional statistical methods with a multitude of temporal and spatial parameters requiring strict statistical correction procedures.

Importantly, electrode-space MVPA revealed better classification accuracy for lMC-tACS, not only at and around the stimulation electrodes FC3 and CP3 but also at left parietal electrodes. It was therefore essential to investigate which specific region was involved using source reconstruction methods. To this end, we analyzed differences in classification accuracies on reconstructed signals from left M1, S1, PMC, SPL, and IPL. We found that classification accuracies in left PMC were significantly stronger comparing lMC-tACS to rCB-tACS/sham for both α and θ in both task and rest signals. This result testifies for aftereffects in premotor but not the primary motor cortex, as the computational model suggested.

## Conclusions

Our results demonstrate a clear broadband aftereffect of 10Hz tACS on the premotor cortex, independent of the state (both task and rest). Specifically, we show that MVPA is a powerful method that enables detection of subtle, single-trial, power changes following 10Hz tACS to the motor cortex. These results have important consequences for therapeutic interventions using tACS since they suggest focal neuroplastic modulation of the motor cortex.

## Acknowledgements

The authors declare no competing financial interests. ET is supported by the DFG grant TZ 85/1-1. We acknowledge support from the University of Leipzig for Open Access Publishing.

