## supplementary materials for "Multivariate pattern analysis reveals location specific aftereffects of 10Hz motor cortex transcranial alternating current stimulation"

1. **Serial reaction time task**

During tACS and at EEG recordings prior and following tACS, participants performed a modified version of the serial reaction time task (SRTT). In each trial, four squares were presented on a grey background in a horizontal array, with each square (from left to right) associated with one of four fingers of the right hand (Supp. Fig. 1A). At stimulus onset, one of the squares turned blue and the rest remained black. Participants were instructed to respond to this blue colored square with the corresponding button, as precisely and quickly as possible. The stimulus remained on the screen until a button press was registered. In case of a wrong button press, the blue colored square turned to red to mark the error. The response-stimulus interval was 500ms (Supp. Fig. 1B). Trials were counted as correct when the appropriate key was pressed within 1000ms after stimulus onset. In case no button was pressed within this time frame, a text appeared on the screen requesting the participants to be faster (“Schneller!”).

The task consisted of three different conditions: simple (SMP), random (RND), and sequence (SEQ). In SMP, stimuli were presented in a simple order of button presses 4-3-2-1-4-3-2-1. In RND, stimuli were presented in a pseudorandom order, generated using Matlab (The Mathworks®, Natick, MA), such that items appeared exactly twice, were not repeated and pairs of consecutive stimuli were followed by some other stimuli, thereby preventing learning by pairwise associations [1]. In SEQ, stimuli were organized in an 8-items-sequence (4-1-4-2-3-1-3-2) also preventing pairwise associations.

**B**

**A**

500ms

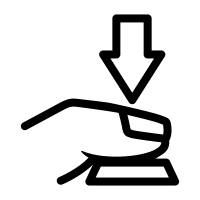

Time

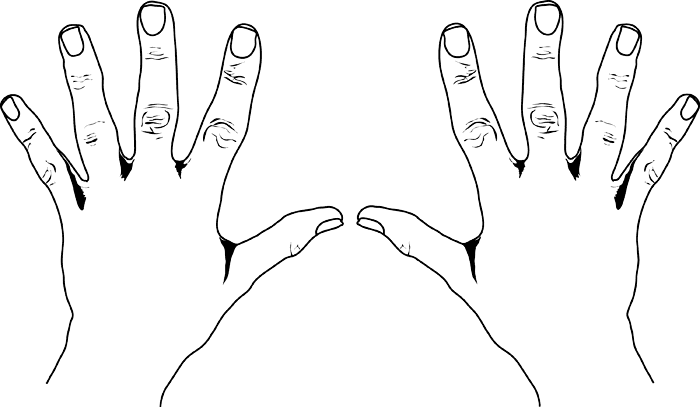

***Supp. Figure 1.*** Serial reaction time task. In each trial, 4 black squares were presented. At stimulus onset, one of the squares turned blue and subjects were instructed to press the button corresponding to the blue square with the respective finger. **B** Task timeline.

During tACS, participants performed a total number of 11 blocks, separated by 20s breaks. There was one SMP block at the beginning or the end of the task (counter-balanced between subjects), and five RND blocks and five SEQ blocks in an alternating order. Each of the SEQ blocks contained 15 repetitions of the 8-element sequence summing to a total of 120 trials per block. The SMP block contained 10 repetitions of the simple sequence summing to a total of 80 trials. Each of the RND blocks contained 80 trials.

During EEG recordings prior tACS, participants performed two RND blocks, and following tACS two RND and two SEQ blocks in alternating order. For the analyses described in the main text only the second RND block in PRE-tACS and the first RND block in POST-tACS were analyzed.

**PRE-tACS**

**tACS**

**POST-tACS**

RND

RND

SEQ

120x4

SEQ

…

SEQ

…

120x3

120

80

SEQ

40

120

40

40

RND

80

80

SEQ

120

RND

80

SMP

RND

RND

RND

2. **Computational modelling of M1 and cerebellar-stimulation locations**

We used the SimNIBS software package (http://simnibs.org/), version 2.1 [2], to simulate the optimal electrode montage for focal left motor cortex (lMC) and right cerebellar (rCB) electric stimulation. The head model, provided by the software package, was created using finite element modeling on T1- and T2-weighted MRI images of an exemplary subject, resulting in a high-resolution tetrahedral head mesh model containing 6 tissue types (grey matter (GM), white matter (WM), cerebrospinal fluid (CSF), skull, skin and eye balls). We set the following standard conductivity values for the 6 tissue types: WM: 0.126 S/m, GM: 0.275 S/m, CSF: 1.654 S/m, skull: 0.01 S/m, eye balls: 0.500 S/m as well as the following conductivity values for the electrode rubber = 29.4 S/m and the electrode gel = 1.0 S/m. All tissues were treated as isotropic. The electrical field E was determined by taking the numerical gradient of the electric potential. For both montages, we used ring-shaped electrodes with 48mm outer diameter, 24mm inner diameter and 3mm thickness. The size and geometry of the electrodes were incorporated into the forward model. The total current injected was 1mA. For lMC-tACS montage, electrodes were placed at EEG locations FC3 and CP3. For rCB-tACS montage, one electrode was placed 1cm below and 3cm right to the inion and the other over right mandibula.

3. **General tACS effects**

None of the subjects reported any adverse effects during or after stimulation. We asked subjects whether they experienced phosphenes, which could be a sign of visual cortex stimulation [3]. Seven out of 25 subjects reported phosphenes during either lMC-tACS (N = 2) or rCB-tACS (N = 5). Three of the seven claimed to see phosphenes during sham. There were no reports on pain or dizziness due to stimulation. During real tACS when asked whether the session was sham, subjects correctly answered “no” in 48% of all real tACS sessions. During sham, 19 out of 25 subjects answered correctly that the session was sham, which might indicate that they were aware of the intervention. However, it seems that the formulation of the question (“was this session sham?”) has led to this large number of subjects correctly identifying the sham session. For example, five out of 25 subjects answered “yes” in all sessions, and 13 out of 25 subjects answered “yes” in two out of three sessions.

4. **Source reconstruction**

To reconstruct the EEG signals in source space we used the linear constrained minimum variance (LCMV) beamforming approach. As a first step we created a head model which was used to estimate the electric field measured by the EEG electrodes. Since individual MRI scans were not available, a standard MRI template was used to construct the boundary element model. To this end, we segmented a template into three tissue types: brain, skull and scalp. Next, we estimated for each tissue type a boundary triangle mesh (brain: 3000 points, skull: 2000 points and scalp: 1000 points). Based on this geometry, a volume conduction model was specified (using standard tissue conduction values) using a Boundary Element Method. For each grid point, we calculated a lead field matrix which was then used to calculate the inverse spatial filter. The inverse spatial filter was calculated across all trials (PRE & POST). Source orientation was optimized by using the orientation of maximum signal power [4], resulting in a reconstructed signal in the alpha and theta frequency bands for each grid point.

5. **Phase-coupling analysis**

To examine the relationship between aftereffects in theta and alpha frequency bands, we measured phase-coupling between theta and alpha oscillations using the phase-locking value [5] defined as follows:

$${PLV}_{\theta,\alpha}=\frac{1}{N}\left| \sum exp(i[\emptyset_{\theta}(t)-\emptyset_{\alpha}(t)]) \right|$$

N is the number of trials and $\emptyset(t)$ is the phase of the theta or alpha oscillation at each time point. POST-PRE phase-locking values (PLV) in each stimulation protocol were compared using the non-parametric Kruskal-Wallis test.

We then tested for PLV differences in POST-PRE tACS, averaged across trials, across the stimulation protocols (lMC-tACS, rCB-tACS and sham) using Kruskal-Wallis test in electrodes FC3, C3, and CP3. P-level threshold was FDR-corrected across the three electrodes.

6. ***No associations between classification accuracies and motor performance***

To explore whether better classification of theta and alpha power post-stimulation may relate to performance changes due to tACS, we correlated F1-values for electrode FC3 and CP3 with reaction-time differences between PRE and POST blocks. We found no evidence to support an effect of motor performance on classification accuracies of theta and alpha power in electrode FC3 and CP3 (data not shown).
